# Epigenetic regulation of genome integrity by a prion-based mechanism

**DOI:** 10.1101/152512

**Authors:** James S. Byers, David M. Garcia, Daniel F. Jarosz

## Abstract

Epigenetic mechanisms mediate diverse gene expression programs in growth and development. Yet whether any can permanently alter the genome is unknown. Here we report a protein-based epigenetic element, a prion, formed by the conserved DNA helicase Mph1/FANCM. [*MPH1*^+^] provides resistance to DNA damage, a gain-of-function trait that requires helicase activity and interactions with other DNA repair proteins. Strikingly, the intrinsically disordered regions of Mph1 and human FANCM that are required for prion phenotypes do not resemble known prions. [*MPH1*^+^] reduces mitotic mutation rates, but promotes meiotic crossovers, driving phenotypic diversification in wild outcrosses. Remarkably, [*MPH1*^+^] is induced by stresses in which the prion is beneficial. Thus, [*MPH1*^+^] fuels a quasi-Lamarckian form of inheritance that promotes survival of the current generation and diversification of the next.

## INTRODUCTION

All organisms must faithfully transmit a genetic blueprint comprising ~10^6^ to ~10^9^ base pairs to the next generation. This challenge is met by an ancient cohort of proteins that orchestrate error-free replication and DNA repair (Whitby, 2010). Many such activities are organized by a helicase known as Mph1 (mutator phenotype 1) in fungi and FANCM (Fanconi Anemia complementation group M) in animals and plants (Crismani et al., 2012; Moldovan and D’Andrea, 2009). Mph1/FANCM also drives diverse biochemical activities including DNA-dependent ATPase function, replication fork reversal, and D loop dissociation (Whitby, 2010). In humans, mutations in FANCM are associated with Bloom’s syndrome and elevated cancer rates, underscoring its central importance in ensuring genome integrity (Alter, 1996; Meetei et al., 2003). A hallmark of Fanconi anemia patients is extreme sensitivity to chemotherapeutics that create DNA inter-strand crosslinks or double strand breaks (DSBs) (Moldovan and D’Andrea, 2009). In nature DSBs arise commonly during sexual reproduction, where they serve as intermediates for meiotic recombination, but vastly exceed the number of crossovers in most organisms (McMahill et al., 2007). Recent studies in *Arabidopsis thaliana* and *Schizosaccharomyces pombe* suggest that, by suppressing crossover events, Mph1/FANCM provides a molecular explanation for this imbalance (Crismani et al., 2012; Lorenz et al., 2012).

Here we report that Mph1 has the capacity to act as protein-based epigenetic element, a prion, that we term [*MPH1*^+^] (brackets denote non-Mendelian meiotic segregation; capitalization denotes dominance). [*MPH1*^+^] is adaptive during genotoxic stress. It also exerts a strong influence on genome fidelity, decreasing mutagenesis in mitotic growth while increasing the frequency with which DSBs are resolved as crossovers during sexual reproduction. [*MPH1*^+^] can be induced by the same stresses to which it provides resistance, providing a molecular mechanism for quasi-Lamarckian inheritance. Our data thus establish [*MPH1*^+^] as an inducible epigenetic state that simultaneously guards the genome against insult during mitotic division and promotes its permanent and heritable diversification during meiosis.

## Methods

### Yeast Techniques

Yeast strains (Table S1) were obtained from stock centers or generously provided by the sources indicated. All strains were stored as glycerol stocks at −80 °C and revived on YPD before testing. Yeast were grown in YPD at 30 °C unless indicated otherwise. Yeast transformation was performed with a standard lithium–acetate protocol (Gietz et al., 1992).

For yeast crossing experiments, MATa and MATα mating types each lacking a unique auxotrophic marker were mixed overnight in YPD. This mixture was then plated on plates selecting for these 2 auxotrophies to select for diploids (usually minus lysine and minus methionine). After 3 days, colonies were picked and grown in 8 mLs pre-sporulation media (8 g Yeast Extract, 3 g Bacto Peptone, 100 g Dextrose, 100 mg adenine sulfate per liter) overnight. The next morning, the cells were washed and resuspended in 3 mLs sporulation media (10 g potassium acetate, 1 g yeast extract, 0.5 g glucose, 0.1 g amino acid add-back per liter). Cells were incubated for 7 days at room temperature and spores were enriched as described previously (Rockmill et al., 1991).

Cytoduction experiments were performed as described previously (Chakrabortee et al., 2016). Briefly, initial BY4742 recipient strains generated by transformation introducing a defective KAR allele (*kar1-15*) that prevents nuclear fusion during mating (Wickner et al., 2006). Strains were made ‘petite’ (incompetent for mitochondrial respiration) by inoculating a single colony in YPD with 0.25% ethidium bromide and growing culture at 30 °C until late exponential/stationary phase (OD_600_ ~ 1). Cultures were diluted 1:1000 into fresh YPD with ethidium bromide and the previous steps were repeated twice. Cultures were plated to single colonies and multiple were tested for respiration incompetence (i.e. no growth on YP-Glycerol). For initial cytoduction into BY4742, donor BY4741 strains harboring [*MPH1*+] and naïve BY4742 *kar1-15* recipient strains were mixed on the surface of a YPD agar plate. These were grown for 24 hours and transferred to media lacking methionine and containing glycerol as a carbon source (SGly-MET). This selects for both BY4742 nuclear markers along with restoration of functional mitochondria via cytoplasmic exchange. After 3-5 days, multiple single colonies were picked and passaged with another round of selection on SD-MET. In parallel, these colonies were confirmed to be haploid by passage on SD-LYS-MET medium. For reverse cytoductions, the new donor strains (this time the successful BY4742 *kar1-15* cytoductants) were mixed with petite naïve BY4741 cells (generated as above) on YPD agar. Cytoductions were repeated as described previously except selecting for BY4741 recipient nuclear markers on glycerol media lacking lysine (SGly-LYS). Multiple cytoductants were picked and tested for the presence of [*MPH1*^+^] phenotypes.

### Phenotypic assays

Biological replicates of each yeast strain (BY4741 MATa haploids) were pre-grown in rich media (YPD). We then diluted these saturated cultures 1:10 in sterile water and then inoculated 1.5 μL into 96-well plates with 150 μL of YPD (SD-CSM was used instead with cisplatin and mitomycin C) per well with the following stressors: phleomycin 5μg/mL; mycophenolic Acid – 20 μM; hydroxyurea – 400 mM; 4-nitroquinoline 1-oxide – 1.2 μM; cisplatin – 8 mM; methyl methanesulfonate – 0.012%; camptothecin –50 μM; doxorubicin – 80 μM; oxolinic acid –100 μM; mitomycin C –1mM. We grew cells at 30 °C in humidified chambers for 96 h and continuously measured growth by OD_600_ in a microplate reader. Timepoints plotted in bar graphs correspond to the point of greatest difference between [*mph1*^-^] and [*MPH1*^+^]: phleomycin – 1225 min; mycophenolic Acid – 755 min; hydroxyurea – 2500 min; 4- nitroquinoline 1-oxide – 2500 min; methyl methanesulfonate – 1600 min; camptothecin – 900 min; doxorubicin – 2000 min; oxolinic acid – 755 min; mitomycin C – 2500 min.

For genetic interaction experiments, phenotyping was performed similarly to before but instead in 384-well plates to generate additional technical replicates for each strain. Briefly, biological replicates of each knockout were pre-grown in rich media (YPD) in 96-well plates. We did not analyze mutants of two additional interacting proteins, Rfa1 and Smc5, which were associated with strong sporulation defects unrelated to the prion (Farmer et al., 2011; Soustelle et al., 2002)). We then diluted these saturated cultures 1:10 in sterile water and then inoculated 5 μL in a 1:4 array into 384-well plates with 45 μL of YPD+4-NQO per well. However, because each genetic knockout was differentially sensitive to genotoxic stress, different concentrations of the chemical were used for each genotype: *rad5*Δ – 2 μM; *rad51*Δ – 1.5 μM; *sgs1*Δ – 1 μM; *srs2*Δ – 2 μM; *mhf1*Δ – 1.5 μM; *mhf2*Δ – 2 μM; *chl1*Δ – 2 μM; *exo1*Δ – 1.5 μM; *mgm101*Δ – 1 μM; *msh2*Δ – 1 μM; *msh6*Δ – 2 μM; *pso2*Δ – 1.5 μM; *slx4*Δ – 2 μM. We grew cells at 30 °C in humidified chambers for 96 h and continuously measured growth by OD_600_ in a microplate reader. The Log2 fold-change values plotted correspond to the highest OD_600_ value reached for each strain (i.e. the carrying capacity). The same concentrations were used for the phenotyping of the corresponding heterozygous, cross-back diploids.

### Microscopy

Microscopy was performed using a Leica inverted fluorescence microscope with a Hammamatsu Orca 4.0 camera. Cells were imaged after growth to exponential phase (OD600 of 0.7) or stationary phase (OD 1.5 or above after 2 days) in a medium that minimizes autofluorescence (per liter in water: 6.7 g yeast nitrogen base without ammonium sulfate, 5 g casamino acids, 20 g glucose). Exposure time was 2 seconds.

### Protein Transformation

Lysate transformations were performed as described previously (Chakrabortee et al., 2016; Tanaka and Weissman, 2006). Briefly, 50 mL cultures of [*MPH1*^+^] strains were grown in YPD for 18 h, pelleted, washed twice with H_2_O and 1 M sorbitol respectively, and then resuspended in 200 μL of SCE buffer (1 M sorbitol, 10 mM EDTA, 10 mM DTT, 100mM sodium citrate, 1 Roche mini-EDTA-free protease inhibitor tablet per 50 mL, pH 5.8) containing 50 units/mL of zymolyase 100T. Cells were incubated for 30 min at 35°C, sonicated on ice for 10 s with a sonic dismembrator at 20% intensity, and cell debris was removed via centrifugation at 10,000g at 4°C X 15 min. Supernatants were digested with 3-fold excess RNAse I and biotinylated DNAse (as determined by units of activity) (Thermo AM1906) for 1 h at 37°C. DNase was subsequently removed by adding saturating quantities of streptavidin-sepharose beads, incubating for 5 min, and bead pelleting via centrifugation (this allows addition of a *URA3*- marked plasmid for selection later). Nuclease digested supernatants were used to transform naïve recipient yeast spheroplasts. Cells were harvested as before, re-suspended in 200 U/mL zymolyase 100T in 1 M sorbitol, and incubated at 35°C for 15 min. Spheroplasts were collected by centrifugation, and washed twice with 1 mL sorbitol and 1 mL STC buffer (1 M sorbitol, 10 mM CaCl2, 10 mM Tris pH 7.5) respectively, and re-suspended in STC buffer using wide-mouthed pipet tips to avoid lysis. Aliquots of spheroplasts were transformed with 50 μL of lysate, 20 μL salmon sperm DNA (2 mg/mL), and 5 μL of a carrier plasmid (*URA3*- and GFP-expressing pAG426-GFP). Spheroplasts were incubated in transformation mix for 30 min at room temperature, collected via centrifugation, and resuspended in 150 μL of SOS-buffer (1 M sorbitol, 7 mM CaCl2, 0.25% yeast extract, 0.5% bacto-peptone). Spheroplasts were recovered at 30°C for 30 min and the entire culture was plated on SD-URA plates and overlaid with warm SD-CSM containing 0.8% agar. After 2-3 days, dozens of Ura+ colonies were picked and re-streaked on SD-URA selective media. The carrier plasmid was subsequently removed by section on 5- FOA and single colonies were tested for the transmission of [*MPH1*^+^]-dependent phenotypes. Infectivity is calculated as percent of transmission divided by the amount of seeded protein used (estimate 77 molecules/cell of Mph1) (Kulak et al., 2014).

### Mutagenesis assays

Yeast strains were grown in multiple biological replicates to saturation in YPD. Then 1 mL was spun down, resuspended in 100 μL H_2_O, and plated on SD-Arg (6.7 g yeast nitrogen base without ammonium sulfate, 5 g casamino acids without arginine, 20 g glucose per liter) + 60 μg/mL Canavanine (forward mutagenesis), YPD + 100 μM Fluconazole, or 60 μg/mL Canavanine and 1 g/L 5- Fluoroorotic acid (GCR mutagenesis). Plates were incubated for 3 days and then CFUs were counted using a colony counter (Synbiosis Acolyte).

### Induced mutagenesis and prion switching assays

Yeast strains (BY4741 or *MDG1::K.lactisURA3* reporter strains) were grown with 3 biological replicates to saturation in YPD with the indicated chemicals for induced mutagenesis frequencies (0.012% MMS, 100 μM Oxolinic Acid) or for [*MPH1*^+^] reporter switching (2.4 μM 4-NQO, 400 μM Camptothecin, 2 mM Cisplatin, 0.024% H_2_O_2,_ 400 mM Hydroxyurea, 0.012% MMS). For starvation conditions in [*MPH1*^+^] reporter switching, yeast strains were grown with 3 biological replicates to saturation in YPD and then washed once with water and resuspended in the following media conditions (no glucose, nitrogen limitation, phosphate limitation, and sporulation media). Then 1 mL was spun down, resuspended in 100 μL H_2_O, and plated on the indicated selective plates: SD-Arg (6.7 g yeast nitrogen base without ammonium sulfate, 5 g casamino acids without arginine, 20 g glucose per liter) + 60 μg/mL Canavanine (forward mutagenesis) or SD-URA (50 mg uracil, 6.7 g yeast nitrogen base without ammonium sulfate, 5 g casamino acids, 20 g glucose per liter) + 1 g/L 5-FOA (reporter switching). Plates were incubated for 3 days and then CFUs were counted using a colony counter (Synbiosis Acolyte). For prion reporter switching, *MDG1* has no defined function in mutagenesis or DNA repair, but is down-regulated in [*MPH1*^+^] cells. Thus, [*mph1*^-^] cells can grow on media lacking uracil, but [*MPH1*^+^] cells cannot. In contrast, [*MPH1*^+^] cells can grow on 5-FOA, whereas [*mph1*^-^] cells cannot.

### Meiotic Recombination Assays

Meiotic reporter strains were generated by amplifying a *K.lactis URA3* cassette off the *pUG72* plasmid (Euroscarf) with primers targeting the marker 50kb upstream of the *his3*Δ locus of BY4741. These PCR products were transformed into [*mph1^-^*], [*MPH1*^+^], and *mph1*Δ strains as described above. A functional *HIS3* marker was also re-integrated back into its endogenous locus via PCR into *BY4742* [*mph1^-^*] wild-type and *mph1*Δ. *BY4741* and *BY4742* strains of the corresponding *MPH1* genotype were crossed to generate His+ Ura+ diploids and then sporulated as described above. Dozens of colonies were picked for each and tested for co-segregation of *URA3* and *HIS3* markers.

For the genetic cross experiments with lab and clinical strains, the [*mph1*^-^] and [*MPH1*^+^] laboratory strains above were crossed to the clinical isolate YJM975 (SGRP). Diploids were sporulated as before and then 96 spores were picked for each. Spores were pinned in quadruplicate onto solid YPD plates with the follower stressors: 39°C, 128 μg/mL fluconazole, 1 mM amphotericin B, 120 μg/mL calcofluor white, 0.01% H_2_O_2_.Yeast were grown at 30°C (except in the case of 39°C heat stress) for 3-4 days and then pictures were taken and colony size analysis was conducted using SGAtools (Wagih et al., 2013). Distributions were normalized to the means of the population.

## RESULTS

### [*MPH1*^+^] is a protein-based genetic element

We previously found that transient overexpression of Mph1 could elicit heritable zinc resistance that had properties consistent with protein-based inheritance (Chakrabortee et al., 2016). These included non-Mendelian segregation of this phenotype in genetic crosses (it was inherited by all meiotic progeny rather than half), and a strong reliance on molecular chaperones (Hsp70 proteins) for propagation from one generation to the next. This chaperone dependence was unusual – most previously known prions depend on Hsp104 to propagate (Shorter and Lindquist, 2005) – and Mph1 lacks the glutamine or asparagine-rich regions typical of nearly all known prions. We therefore investigated whether this Mph1-dependent epigenetic state was a *bona fide* protein-based genetic element.

Prion acquisition commonly elicits heritable changes in the localization of causal protein(s) (Derkatch et al., 2001). We investigated whether this was true for the Mph1-dependent epigenetic state, taking advantage of the fact that prions are dominant (Shorter and Lindquist, 2005). We crossed a yeast strain expressing an endogenous *MPH1-YFP* fusion to a strain harboring the Mph1-dependent epigenetic state, and to genetically identical (isogenic) naïve cells as a control. There was no apparent difference between these groups in exponentially growing cells. However, in stationary phase cells, Mph1-YFP foci were evident in diploids harboring the Mph1-dependent epigenetic state, but not in isogenic naïve diploids (Fig. S1).

We next tested whether the Mph1-dependent epigenetic state could be transmitted by cytoplasmic mixing without transfer of any nuclear material, another hallmark of prion biology (Wickner et al., 2006). To do so, we performed ‘cytoduction’ experiments with *kar1-15* mutants in which nuclei do not fuse during mating (Fig. S2A; see methods). The Mph1-dependent epigenetic state was robustly transferred to naïve recipient cells through such ‘cytoduction’ experiments (Fig. S2B). These data thus establish that the phenotype did not arise from genetic mutation.

Finally, we performed a protein transformation as the ‘gold-standard’ test for prion-based inheritance. We generated nuclease-digested lysates from cells harboring the Mph1-dependent epigenetic state and from isogenic naïve cells. We used these lysates to transform naïve spheroplasts (yeast lacking a cell wall), including a carrier plasmid harboring a *URA3* marker to enrich for cells that were competent to uptake molecules from their external milieu (Fig. S2C; see methods). We picked dozens of Ura+ colonies, propagated them on 5-FOA to select for loss of the carrier plasmid, and characterized the phenotypes of the resulting cells. Strikingly, over 40% of them also acquired the Mph1-dependent epigenetic state (scored by resistance to ZnSO_4_; Fig. 1A; *p*=1.6x10^-4^ by t-test). When benchmarked against the low natural abundance of Mph1 (~80 molecules per cell (Kulak et al., 2014)), this frequency of transmission is substantially more efficient than for other prions that have been tested (Tanaka and Weissman, 2006). We conclude that the Mph1-dependent epigenetic state is a *bona fide* protein-based genetic element – a prion – which we hereafter refer to as [*MPH1*^+^].

**Figure 1.**
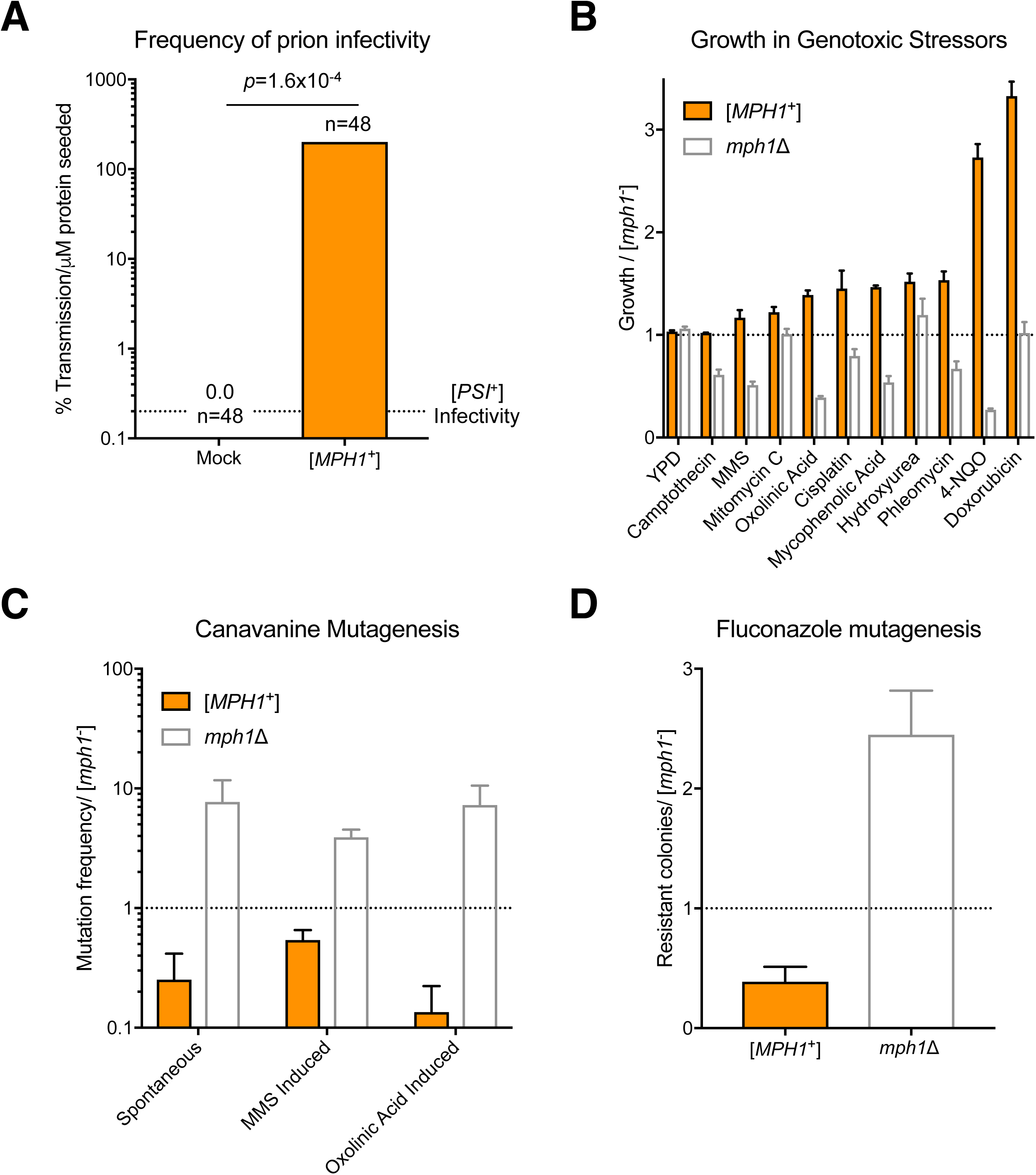
[*MPH1*^+^] is a prion that protects against DNA damage mutagenesis. (A) “Frequency of prion infectivity” for protein transformation comparing [*mph1*^-^] lysates (Mock) and [*MPH1*^+^]-harboring lysates. Infectivity is calculated as described in methods. [*MPH1*^+^] was scored by resistance to ZnSO_4_ (Chakrabortee et al., 2016). Infectivity from assembled [*PSI*^+^] is displayed as a dotted horizontal line (Tanaka and Weissman, 2006). (B) Growth of [*MPH1*^+^] and *mph1*Δ cells in genotoxic stress relative to naïve [*mph1*^-^] cells. Error bars represent SEM from three or more biological replicates. (C-D) Spontaneous and induced mutation frequencies of [*MPH1*^+^] and *mph1*Δ cells, relative to naïve [*mph1^-^*], after growth in YPD (spontaneous) or after exposure to mutagenic agents (ex. 0.012% MMS or 100 μM oxolinic acid) (see methods for details). Naïve mutation frequencies increased dramatically after exposure to MMS and oxolinic acid (5x10^-4^ and 2x10^-5^ mutants per CFU respectively). Error bars represent SEM from 3 independent experiments.

### [*MPH1*^+^] promotes DNA damage tolerance and genome stability

Because Mph1 plays a central role in genome integrity we tested the ability of [*MPH1*^+^] cells to withstand DNA damage. The prion had no effect on growth in rich medium alone. We next exposed [*mph1*^-^], [*MPH1*^+^], and *mph1*Δ cells to a battery of genotoxic insults, including replication stressors (hydroxyurea, mycophenolic acid), intercalating agents (doxorubicin), lesion inducers (4-NQO), alkylating agents (MMS), topoisomerase inhibitors (oxolinic acid), DNA break inducers (phleomycin), and crosslinkers (cisplatin, camptothecin, mitomycin C). In most of these conditions, cells harboring [*MPH1*^+^] grew better than isogenic [*mph1*^-^] cells, and the prion was never maladaptive (Fig. 1B). In contrast, *mph1*Δ cells were hypersensitive to most genotoxic stressors. Many amyloid-based prions act by sequestering their constituent protein into aggregates. Thus their phenotypes often mimic the corresponding genetic loss-of-function. Our data indicate that [*MPH1*^+^] is capable of driving gain-of-function phenotypes.

Mph1 overexpression strongly increases the frequency of gross chromosomal rearrangements (GCR) and, conversely, *mph1*Δ decreases GCR formation (Banerjee et al., 2008). We observed no influence of [*MPH1*^+^] on GCRs using the same reporter strains employed in those studies (*p*=0.40 by Student’s t-test, Fig. S3) however, establishing that the prion state is distinct from a simple increase in Mph1 activity. We also examined mutagenesis using an assay in which loss-of-function mutations in the *CAN1* gene render cells resistant to the toxic arginine analog canavanine. As others have reported (Scheller et al., 2000), *mph1*Δ cells had an approximately tenfold increased spontaneous mutagenesis frequency compared to wild-type cells. In contrast, [*MPH1*^+^] cells had a three-fold *decreased* frequency of spontaneous mutation compared to isogenic [*mph1*^-^] cells (Fig. 1C), again consistent with [*MPH1*^+^] driving gains-of-function. We observed the same anti-mutator phenotype when scoring resistance to the antifungal drug fluconazole (Fig. 1D), establishing that these relationships were not unique to the forward mutagenesis assay employed. We also observed decreased frequencies of induced mutagenesis in [*MPH1*^+^] cells (e.g. with MMS and oxolinic acid; Fig. 1C).

One possible explanation for the decreased mutation frequency in [*MPH1*^+^] cells would be preferential engagement of error-free repair pathways, which often involve homologous recombination. Indeed, Mph1 has been implicated in pathway choice decisions in multiple organisms (Huang et al., 2013; Xue et al., 2015b) and [*MPH1*^+^] provides resistance to agents that induce DSBs (*e.g*. phleomycin; Fig. 1B), which serve as precursors to HR. Using a simple assay for integration of a *HIS3* marker at its endogenous locus (see methods), we found that [*MPH1*^+^] drives a robust increase in HR (5.3-fold; *p*=0.003 by t-test; Fig. S4), providing a possible explanation for the reduced mutagenesis frequency. Collectively, our data establish that [*MPH1*^+^] enhances survival and preserves genome integrity during wide range of genotoxic insults.

### [*MPH1*^+^] phenotypes require helicase function

The gains-of-function in [*MPH1*^+^] cells led us to investigate whether Mph1’s catalytic activity was required for prion-dependent phenotypes. To test this, we employed a well-characterized inactivating point mutation in Mph1’s active site (*mph1-Q603D*, (Chen et al., 2009)). We crossed isogenic [*MPH1*^+^] and [*mph1*^-^] cells to a naïve strain harboring *mph1-Q603D*, selected diploids, sporulated them, and then examined meiotic progeny that harbored the *mph1-Q603D* alleles (Fig. 2A).

**Figure 2.**
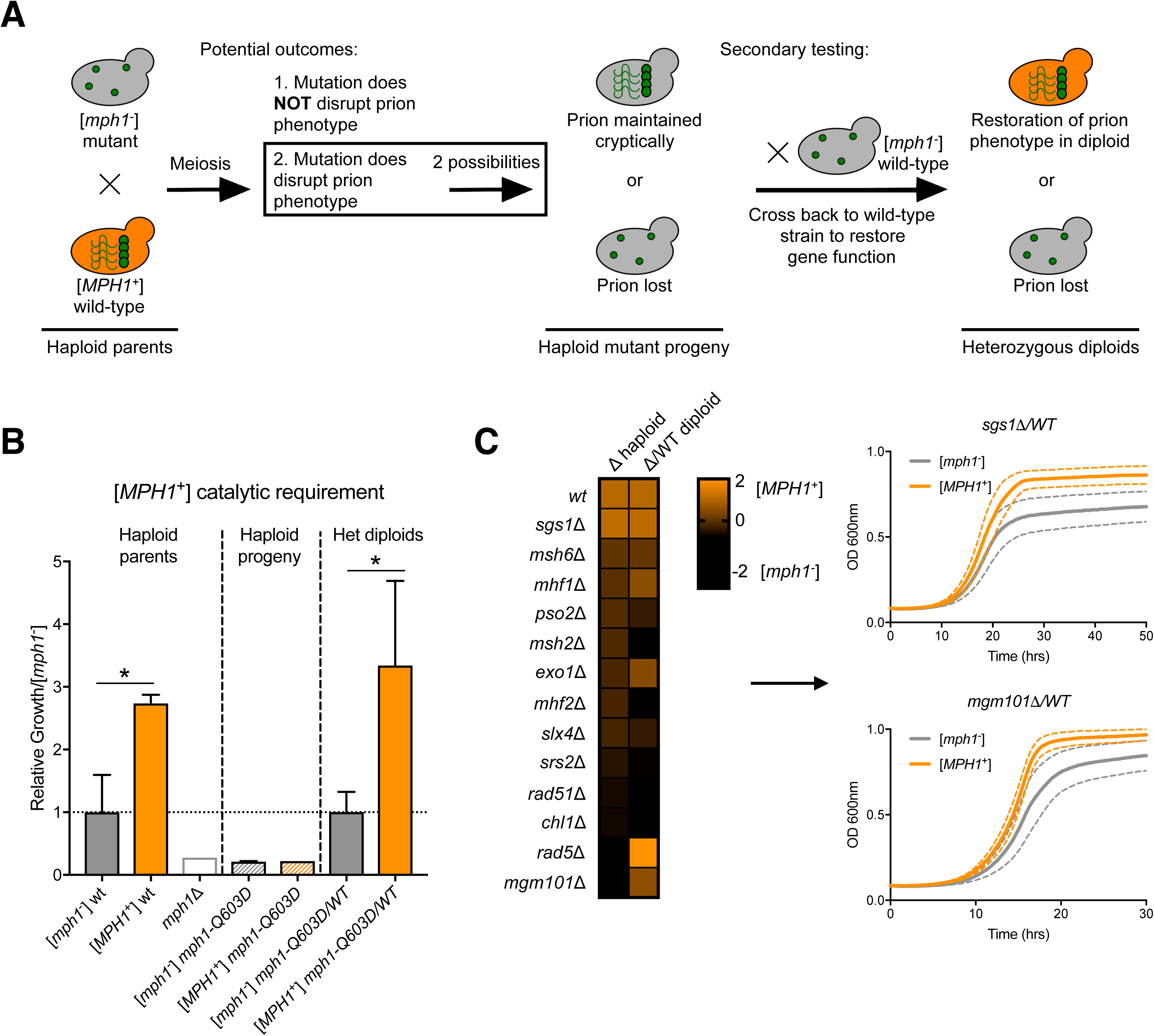
[*MPH1*^+^] requires helicase activity and FA pathway components. (A) Experimental schema for crossing [*MPH1*^+^] into mutant strains to determine whether [*MPH1*^+^]’s catalytic activity or interacting proteins are required to manifest prion phenotypes (in haploid progeny) and/or propagate the prion. (B) Haploid parents: growth of wild-type [*mph1^-^*] and [*MPH1*^+^] in the DNA damage stressor 4-NQO (1.2 μM). Haploid mutant progeny: growth of [*mph1^-^*] and [*MPH1*^+^] strains harboring a single catalytically dead point mutation (*mph1-Q603D)* in the *MPH1* gene in 1.2 μM 4-NQO. Heterozygous diploids: growth of [*mph1^-^*] and [*MPH1*^+^] catalytically dead mutants in 1.2 μM 4-NQO after crossing back to a wild-type strain. Error bars represent SEM determined from 6 biological replicates and are present on all plots. (C) Left – Δ haploid: heatmap of Log2-transformed fold-changes in growth during DNA damage (4-NQO; see methods) of haploid progeny harboring deletions of Mph1-associated proteins after crossing with an [*MPH1*^+^] strain. Data are normalized to isogenic [*mph1^-^*] controls. Δ/WT diploid: data as above, but for heterozygous diploids in which gene function was restored. Right – representative growth curves of cross-back, heterozygous diploids in which the [*MPH1*^+^] phenotype was maintained or restored. Error bars represent SEM determined from 6 biological replicates.

As expected, [*MPH1*^+^] progeny harboring the wild-type allele were more resistant to 4-NQO than matched [*mph1*^-^] progeny (*p*=0.014 by t-test, Fig. 2B). In contrast, [*MPH1*^+^] progeny harboring the *mph1*- *Q603D* allele were as sensitive to 4-NQO as [*mph1*^-^] progeny harboring the catalytically inactive variant (*p*=0.21, Fig. 2B). Thus, Mph1 activity is required to produce [*MPH1*^+^] phenotypes. We tested whether Mph1 catalytic function was also required for prion propagation (Fig. 2A). To do so we crossed meiotic progeny (arising from the previous cross) harboring the *mph1*-*Q603D* allele to naïve, wild-type strains to create heterozygous diploids in which Mph1 function was restored. Prion-dependent resistance to 4- NQO re-emerged in these strains (*p*= 0.047 by t-test, Fig. 2B), establishing that the helicase function of Mph1 is required to produce DNA damage resistance, but not to propagate the prion itself.

### [*MPH1*^+^] phenotypes require FA pathway components

As a scaffold, Mph1 interacts with many factors involved in both error-free and error-prone DNA repair (Daee et al., 2012; Ward et al., 2012; Xue et al., 2015a). We applied classical epistasis to examine how these interactions influence [*MPH1*^+^]-dependent phenotypes, capitalizing on the fact that [*MPH1*^+^] is transmitted to all progeny of meiosis (Chakrabortee et al., 2016). We crossed isogenic [*MPH1*^+^] and [*mph1*^-^] cells to naïve strains harboring deletions of thirteen previously reported Mph1-interacting genes, related helicases, proteins involved in inter-strand crosslink (ICL) repair, and FA pathway components. We then sporulated these diploids and selected six [*MPH1*^+^] and [*mph1*^-^] progeny harboring each gene deletion. Finally, we exposed these strains to genotoxic stress and compared how [*MPH1*^+^] affected survival in the context of each gene deletion.

We observed strong genetic interactions between [*MPH1*^+^] and components of the FA pathway. This network has been best characterized in the context of ICL repair, where Mph1 scaffolds Msh2/Msh6 to recruit Exo1 and digest the ICL-harboring oligonucleotide. Mhf1/Mhf2 stabilizes Mph1 complexes on chromatin at stalled replication forks and Chl1 (the yeast FancJ homolog) and Slx4 (the yeast FancP homolog) mediate downstream gap re-filling and fork reset (Daee et al., 2012). Loss of each of these FA factors eliminated [*MPH1*^+^]-dependent resistance to genotoxic stress (Fig. 2C).

Deletion of two genes encoding proteins related to Mph1/ICL repair, Pso2 (a nuclease and member of an orthogonal ICL epistasis group) (Ward et al., 2012) and Srs2 (the yeast RTEL1 helicase homolog), abolished the [*MPH1*^+^] phenotype, even though they are not linked to the FA pathway (Daee et al., 2012). These epistasis patterns establish that the phenotypic effects of [*MPH1*^+^] require engagement of most yeast Fanconi proteins (Ward et al., 2012), as well as other factors that are thought to function in independent, parallel repair pathways (Fig. 2C), raising the possibility that [*MPH1*^+^] exerts its phenotypes by re-wiring crosstalk among DNA repair factors.

We next tested whether these Mph1 interactions were required for prion propagation. To do so we crossed the haploid deletion strains in which [*MPH1*^+^]-dependent phenotypes had disappeared to naïve wild-type strains, creating heterozygous diploids in which gene function was restored. Strikingly, resistance to genotoxic stress re-emerged in only 3 strains: *exo1*Δ*/EXO1*, *rad5*Δ*/RAD5*, and *mgm101*Δ*/MGM101* (Fig. 2C). Thus, the remaining proteins (Rad51, Srs2, Mhf1, Mhf2, Chl1, Msh2, Msh6, Pso2, Slx4), which include most of the FA pathway, are required not only to manifest [*MPH1*^+^] phenotypes, but also to propagate the prion.

Genetic requirements for mutagenesis and survival in DNA damage can differ. We therefore investigated how the loss of these same genes affected the anti-mutator phenotype of [*MPH1*^+^]. (This was only possible in haploids because most canavanine resistant mutants are recessive.) Cells lacking Mgm101, Pso2, and Rad51 inherited anti-mutator phenotypes from an [*MPH1*^+^] parent (Table S3), but those lacking Rad5, Msh2, Msh6, or Mhf1 did not inherit an anti-mutator phenotype. Strikingly, [*MPH1*^+^] led to large *increases* in mutation frequency in strains lacking Slx4 and Mhf2. Mhf2 stabilizes aberrant DNA structures, allowing helicases such as Mph1 to remodel them. Slx4 is a structure-specific endonuclease that also acts on aberrant DNA structures (particularly branched substrates) (Fricke and Brill, 2003). Slx4 has overlapping specificity with another DNA helicase, Sgs1 (BLM in humans), which interacts with topoisomerase 3 to process stalled replication forks (Fricke and Brill, 2003). As with Slx4, loss of Sgs1 also led to a strong mutator phenotype in the presence of [*MPH1*^+^]. Our data thus strongly suggest that the anti-mutator phenotype in [*MPH1*^+^] cells depends on Slx4, Sgs1, and Mhf2 activities, and likely their ability to resolve aberrant DNA intermediates in a non-mutagenic fashion.

### Prion-dependent toxicity of Mph1 and FANCM overexpression

A hallmark of many prion proteins is that they can be toxic when overexpressed in matched [*PRION*^+^] strains, but not in naïve cells (Fig. 3A). This toxicity typically requires specific protein domains that are important for prion propagation (Douglas et al., 2008). Mph1 overexpression from a constitutive promoter was indeed significantly more toxic in [*MPH1*^+^] cells than in [*mph1*^-^] cells (*p*=0.04 by t-test, Fig. 3B). We used this growth impairment as a tool to dissect which domains of Mph1 fuel its prion-like properties. Mph1 does not harbor N/Q-rich prion-like domains (Fig. 3C). However, it does contain multiple, large, extremely disordered regions outside of its helicase domain (Jones and Cozzetto, 2015) (Fig. 3D), particularly at the C-terminus. This pattern of disorder is highly conserved in the human ortholog FANCM (Fig. 3D), and truncation of the disordered C-terminal region is the most frequent FANCM variant observed in human cancers (Cerami et al., 2012). Notably, Mph1 variants lacking its large C-terminal disordered region did not exert prion dependent toxicity (Fig. 3B), despite similar levels of expression (Fig. S5), establishing the importance of this disordered region in prion-dependent phenotypes.

**Figure 3.**
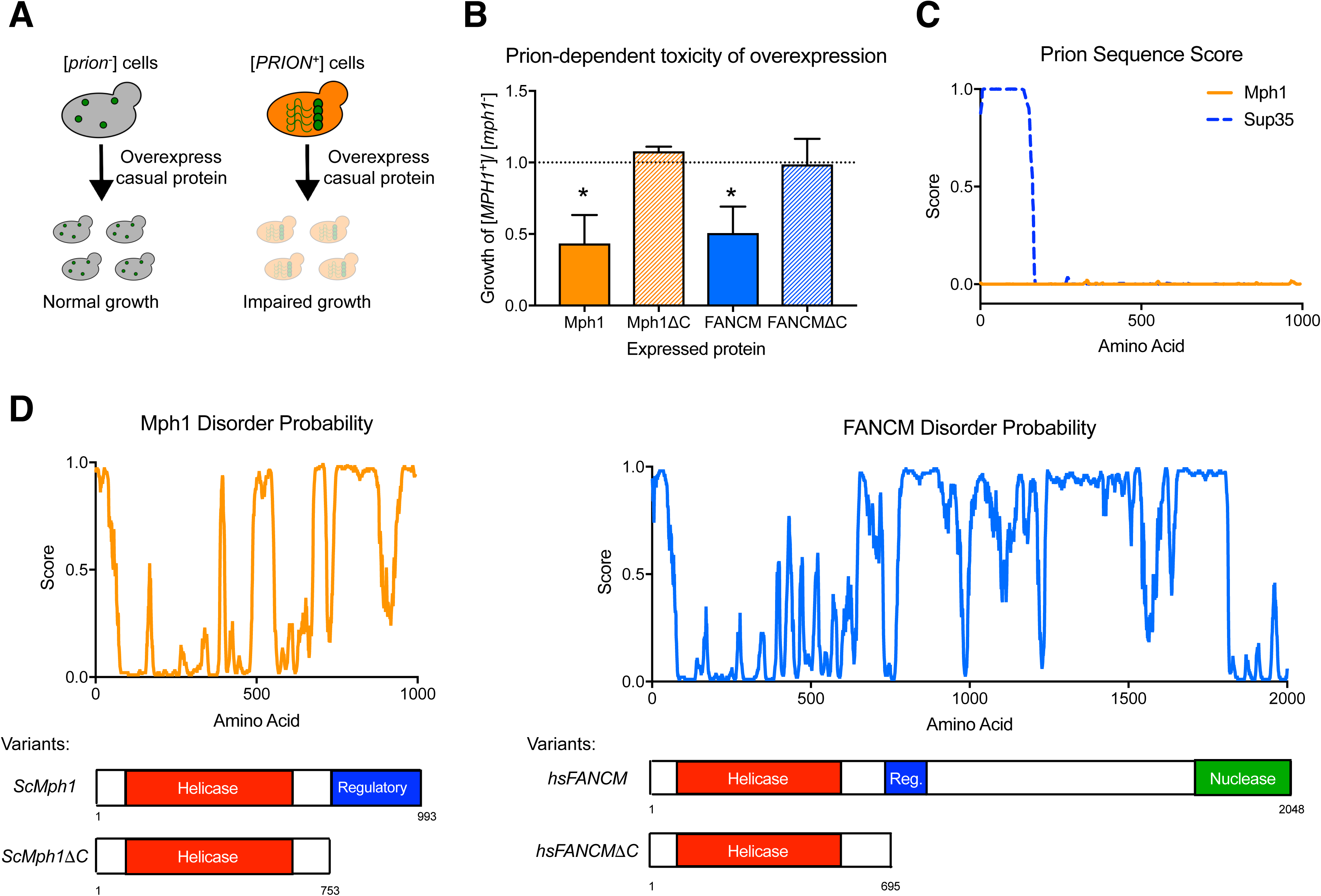
Prion-like behavior and intrinsic disorder is conserved in human FANCM. (A) Schematic depicting the toxicity of casual protein overexpression in the presence of the corresponding prion. (B) Growth of [*MPH1*^+^] strains harboring plasmids expressing FANCM and Mph1 variants. Values are normalized to corresponding [*mph1*^-^] strains harboring the same plasmids. Error bars represent SEM from six biological replicates. (C) Scores from an algorithm that predicts prion-like sequences using a Hidden Markov model (PLAAC; (Lancaster et al., 2014)) for both Mph1 and Sup35. (D) Top: Intrinsic disorder predictions (Disopred3; (Jones and Cozzetto, 2015)) for Mph1 and FANCM. Bottom: Variants and domain architecture of Mph1 and FANCM used in this assay.

We also transformed [*mph1*^-^] cells and [*MPH1*^+^] cells with plasmids encoding full-length FANCM and a variant lacking its disordered C-terminus (FANCMΔC) under the control of the same constitutive promoter. Expression of full-length FANCM was also more toxic in [*MPH1*^+^] cells than in isogenic [*mph1*^-^] cells (*p*=0.036 by t-test, Fig. 3B). In contrast, FANCMΔC did not affect growth of either [*mph1*^-^] or [*MPH1*^+^] cells (*p*=0.47 by t-test, Fig. 3B). Thus, despite its limited sequence identity, human FANCM appears to interact with yeast [*MPH1*^+^] to exert a toxicity that depends upon both its disordered C-terminus and the presence of the prion.

### Induction by environmental stress

It has been suggested that prions might drive a quasi-Lamarckian form of inheritance (Halfmann et al., 2010; True and Lindquist, 2000), fueling heritable and adaptive phenotypic changes in response to transient environmental stressors (Holmes et al., 2013). Many prions can be modestly induced by perturbations that disrupt protein homeostasis (Jarosz et al., 2010). Yet only two, [*GAR*^+^] (Jarosz et al., 2014) and [*MOD*^+^] (Suzuki et al., 2012), are known to be induced by the same stresses to which they provide resistance. We tested whether any of the DNA damaging stresses in which [*MPH1*^+^] provides a benefit might also elicit its appearance, as would be expected for a Lamarckian epigenetic element. We did so using an *MDG1*::*URA3* reporter that we previously established provides a readout of the [*MPH1*^+^] prion state (Chakrabortee et al., 2016) (Fig. 4A).

**Figure 4.**
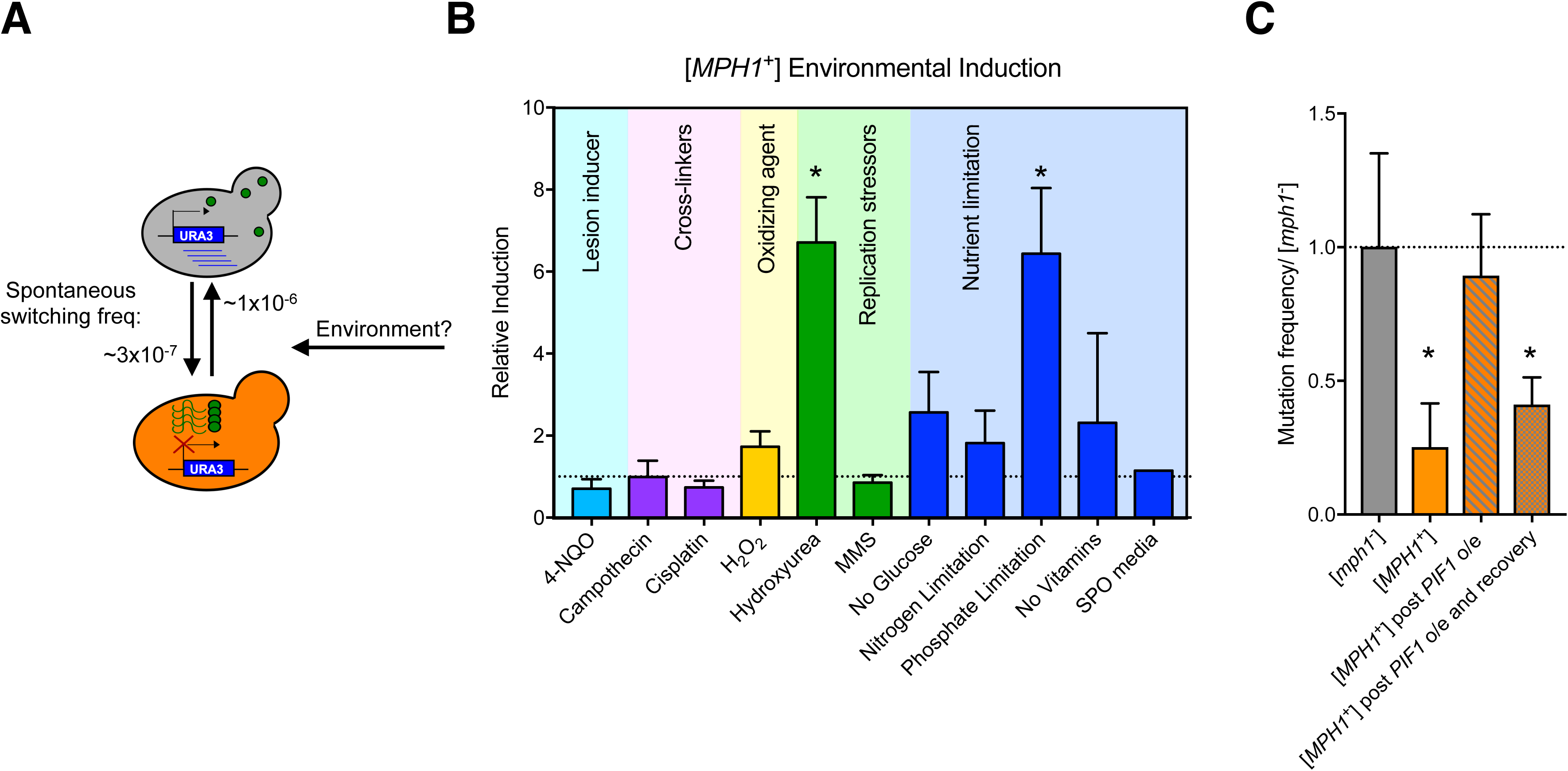
[*MPH1*^+^] is a quasi-Lamarckian element. (A) Diagram of spontaneous switching from [*mph1*^-^] to [*MPH1*^+^]. In this experiment, naïve reporter cells were treated with indicated stressors for 12 hours and assayed for modulated switching frequencies to [*MPH1*^+^]. (B) Induction of [*MPH1*^+^] compared to control in various conditions. Error bars represent SEM from 3 biological replicates. (C) [*MPH1*^+^]- dependent changes in mutation frequency following a transient *PIF1* overexpression to eliminate G-quadruplex regions (Paeschke et al., 2013) and subsequent recovery after ~70 generations in rich media. Values normalized to corresponding [*mph1*^-^] strain. Error bars represent SEM from 8 biological replicates.

Exposure to hydroxyurea increased the frequency of [*MPH1*^+^] acquisition seven-fold in these experiments (p=0.035 by t-test; Fig. 4B). We observed no such effect for many other DNA damaging agents in which [*MPH1*^+^] provides an equivalent or greater adaptive advantage. These data establish that the increased frequency of [*MPH1*^+^] did not merely arise from selection for the prion. Starvation for another nutrient, phosphate, also strongly induced [*MPH1*^+^] (6.4-fold, p=0.041 by t-test). Both hydroxyurea and phosphate starvation have been linked to replication stress via depletion of deoxynucleotide pools (Young et al., 1967). Thus, [*MPH1*^+^] can be induced not just by overexpression, as we did artificially, but also by a specific replication stressor to which it provides resistance, establishing that this prion can act as a quasi-Lamarckian element of inheritance.

Because replication stresses acted as a trigger for [*MPH1*^+^], we wondered whether intermediate structures in DNA repair processes might be important for templating and/or propagating the prion. Mph1 binds to specific DNA fork structures to assemble larger complexes at sites of damage (Xue et al., 2015b). Furthermore, other types of nucleic acids have been hypothesized to act as scaffolds for prion-like aggregation in amyloid bodies (Audas et al., 2016). Constitutive overexpression of Mph1 also leads to a mutator phenotype (Banerjee et al., 2008) that depends upon inactivation of another specialized DNA helicase, Pif1, which efficiently unwinds a type of DNA secondary structure known as G-quadruplexes (Paeschke et al., 2013). These guanine repeats form a stable planar structure through hydrogen bonding, which impedes their traversal by the replication machinery. We *transiently* overexpressed *PIF1* for ~25 generations in [*MPH1*^+^] and [*mph1*^-^] cells and investigated whether they maintained [*MPH1*^+^]-dependent decreases in mutagenesis. Immediately following *PIF1* overexpression, [*MPH1*^+^]-dependent decreases in mutagenesis were completely abolished (*p*=0.40 by t-test, Fig. 4C.). However, after we removed the *PIF1* plasmid and propagated the strains 3 times on standard rich medium (YPD) the [*MPH1*^+^]-dependent mutagenic phenotypes re-emerged (*p*=0.027 by t-test, Fig. 4C). These results indicate that specific DNA structures, such as G-quadruplex regions or perhaps other Pif1 substrates, may play a central role in the manifestation of the [*MPH1*^+^] phenotype, but are not required for propagation of the prion.

### [*MPH1*^+^] increases genetic and phenotypic diversification during meiosis

Experiments in *Arabidopsis thaliana* (Crismani et al., 2012) and *Schizosaccharomyces pombe* (Lorenz et al., 2012) have identified Mph1/FANCM as the strongest known inhibitor of crossover formation during meiosis. We therefore tested whether [*MPH1*^+^] exerted any influence on meiotic crossovers in *S*. *cerevisiae*. To do so we constructed a yeast strain harboring a functional *URA3* cassette 50kb upstream of a non-functional *his3*Δ*1* mutation, enabling us to examine co-segregation of these linked genetic markers (Fig. 5A). We mated this strain, which could grow on media lacking uracil but not on media lacking histidine, to isogenic [*MPH1*^+^] and [*mph1*^-^] strains, which harbored a non-functional *ura3*Δ*0*, but an intact *HIS3* locus. We then selected diploids and induced meiosis. There was no difference in sporulation efficiency between [*mph1*^-^] and [*MPH1*^+^] harboring cells (*p*=0.45 by Student’s t-test, Fig. S6). After 10 days, we isolated meiotic progeny and examined the frequency of His+ Ura+ recombinant spores derived from both [*MPH1*^+^] and [*mph1*^-^] parents. Recombinant phenotypes (*i.e*. those in which re-assortment of linked parental markers occurred) were substantially more common in meiotic progeny derived from [*MPH1*^+^] parents compared to those derived from [*mph1*^-^] parents (4.4-fold; *P* = 0.02 by T-test; Fig. 5B), establishing that the prion fundamentally altered the degree of linkage the cross.

**Figure 5.**
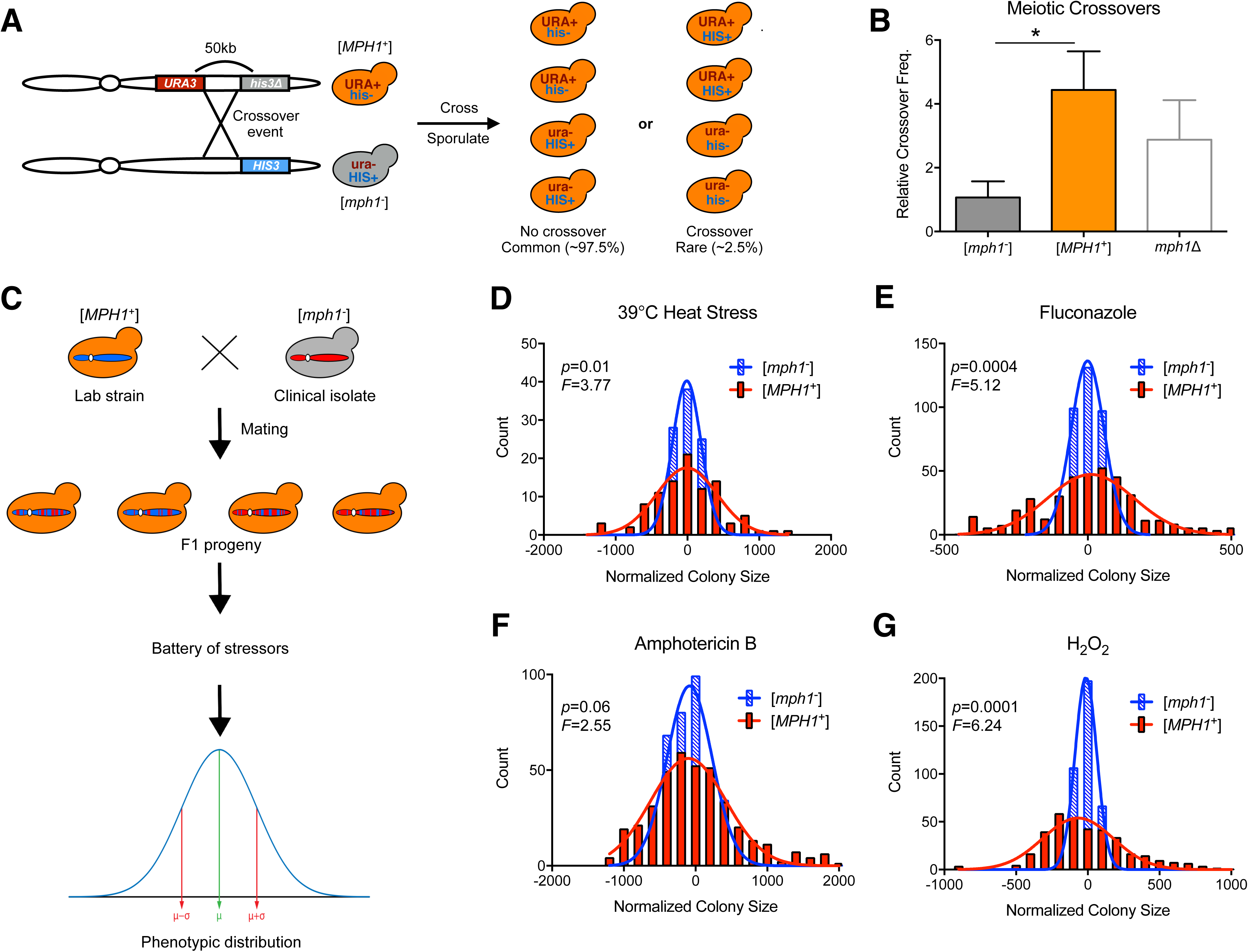
[*MPH1*^+^] increases phenotypic diversification in the progeny of meiosis. (A) Experimental schema for measuring linkage between 2 auxotrophic markers in [*mph1*^-^] and [*MPH1*^+^] strains. (B) Relative frequencies of His+ Ura+ meiotic recombinant progeny in [*mph1*^-^], [*MPH1*^+^], *mph1*Δ strains compared to [*mph1*^-^]. Error bars represent SEM from 4 biological replicates. (C) Experimental schema for determining [*MPH1*^+^]-dependent phenotypic diversification following a cross between lab strains (with or without [*MPH1*^+^]) and a recently evolved clinical pathogen. (D-G) Histograms of normalized spore colony sizes from these crosses (calculated using SGAtools (Wagih et al., 2013)) in 4 clinically relevant stressors. Histograms were fit to a Gaussian distribution to generate curves.

*S. cerevisiae* naturally produces many crossovers per chromosome, a feature that has motivated its use as a genetic model organism. As a consequence, its linkage blocks are small and most polymorphisms within them are thought to be passenger mutations rather than causal variants. We nonetheless investigated whether [*MPH1*^+^] might spark increased phenotypic variation in natural *S*. *cerevisiae* outcrosses. We mated isogenic [*mph1*^-^] and [*MPH1*^+^] laboratory strains to a sequenced clinical strain isolated from an Italian patient (YJM975 (Strope et al., 2015)). As a frame of reference, the genetic divergence between open reading frames in these strains (0.5%) is only slightly greater than that between human individuals. We isolated diploids from these matings, induced meiosis, and isolated spores (Fig. 5C; see SI). We confirmed that these spores were *bona fide* meiotic recombinants based on mating type tests and then exposed these progeny to physiological stressors relevant to the clinical niche: heat stress, antifungal drugs, and oxidative stress, measuring colony size as a proxy for growth. The presence of [*MPH1*^+^] significantly increased the phenotypic variation in these cells (Fig. 5D-G).

To test whether this increased phenotypic diversity arose from the enhanced re-assortment of genetic information during meiosis, or rather due to some other effect of [*MPH1*^+^] (*e.g*. decreased mutation rate, phenotypic capacitance, etc.), we also examined phenotypic variation produced by the prion in the parental strains. We transferred [*MPH1*^+^] to each parent by cytoduction (see SI for experimental details) and examined the variance in phenotype across the same stressors that we used to examine the meiotic progeny. The variance in phenotype imparted by [*MPH1*^+^] was much smaller in each parent than it was in the meiotic progeny (Fig. 5D-G, Fig. S7) and curing of the prion did not eliminate these traits (Fig. S9), establishing that the prion itself did not increase phenotypic variance in a static genetic background. Thus, even given a restricted degree of parental genetic diversity and the high baseline recombination rate of *S. cerevisiae*, [*MPH1*^+^] can sharply increase heritable phenotypic diversification during meiosis.

Changes in protein homeostasis have previously been shown to modulate genotype to phenotype relationships (Jarosz et al., 2010). To confirm that [*MPH1*^+^] was not acting as a phenotypic capacitor and influencing manifestation of the observed phenotypes in the new genetic backgrounds, we picked multiple meiotic progeny and “cured” them of [*MPH1*^+^] via transient inhibition of the molecular chaperone Hsp70 (Chakrabortee et al., 2016). Outlier progeny both resistant and sensitive to heat stress were transformed with a plasmid containing a dominant negative variant of Hsp70 (Ssa1-K69M) and propagated for ~75 generations on selective media to eliminate inheritance of the prion (as has been described previously). Then the plasmid was eliminated and normal Hsp70 function was restored for an additional ~25 generations. The strains were then arrayed onto a plate in a 10-fold dilution series and allowed to grow up for 3 days (Fig. S8). While [*MPH1*^+^]-dependent phenotypes (e.g. zinc resistance) were curable in this experiment, phenotypes that arose uniquely in the meiotic progeny (e.g. resistance to heat stress) were not.

## Discussion

To survive in dynamic, fluctuating environments, organisms must acquire new heritable traits. However, a multitude of mechanisms that safeguard DNA replication often create a phenotypic ‘lock in’, limiting the source of biological novelty to relatively modest changes in the genetic code. Dynamic, environmentally regulated signaling networks offer one solution to this problem. Epigenetic ‘bet-hedging’ mechanisms may provide another (Halfmann et al., 2010; Lancaster and Masel, 2009; True and Lindquist, 2000). Such systems increase phenotypic variation, creating sub-populations that express different traits than the majority. In fluctuating environments, these new traits might enable survival of the population when it would otherwise have perished. However, the evolutionary value of most bet-hedging systems, including those driven by prions, depends upon the constant presence of the causal element. Mechanisms of this type have been implicated in the interpretation of genetic information (True and Lindquist, 2000), but none are known to permanently alter the genome. Our data establish that one such element, the prion [*MPH1*^+^], has the power to do so. This prion is highly transmissible and can be induced in environments where it is adaptive, providing a robust mechanism for Lamarckian inheritance that controls fundamental decisions in DNA damage tolerance, mutagenesis, and meiotic recombination.

Perhaps the greatest force driving genetic diversification in eukaryotes is sexual reproduction. Re-assortment of alleles in meiosis ensures that every genome is fundamentally new. But within this genomic patchwork, linkage blocks can be found in which multiple polymorphisms are inherited in *cis*. As an epistemological tool, geneticists have long assumed that individual, ‘driver’ polymorphisms are linked to many other ‘passenger’ mutations that have no influence on phenotype. Yet evidence from fine mapping studies of individual quantitative trait loci in *S. cerevisiae* (Steinmetz et al., 2002) and metazoans (Mackay et al., 2009) alike suggest that multiple causal alleles can often occur within a single linked genetic locus. The increased phenotypic diversity that we observed in genetic crosses with [*MPH1*^+^] parents suggest, even in crosses with limited genetic diversity and in an organism with small haplotype blocks, that alleles impacting the same phenotype can commonly be linked. This genetic architecture may allow complex traits to persist in a greater number of meiotic progeny, which provides theoretical adaptive advantages. But it also limits meiotic re-assortment of the linked alleles. The [*MPH1*^+^] prion provides a molecular mechanism through which this fundamental decision – whether to couple or separate bits of genetic information as they are broadcast to the next generation – can be reset. The phenotypic consequences of such re-assortment, at least in the context of traits relevant to the clinical niche that we tested, are substantially more adaptive than would be expected from random mutagenesis (where >95% of mutations are detrimental (Eyre-Walker and Keightley, 2007)).

Many prions have long been assumed to be non-functional assemblies of a homogenous protein. However, the fact that the [*GAR*^+^] prion is composed of multiple proteins (Jarosz et al., 2014) suggests that some such elements might function as larger complexes. Indeed, the gain-of-function phenotypes driven by [*MPH1*^+^], as well as inheritance from one generation to the next, appear to be dependent on other interacting DNA repair factors (especially those in the yeast FA pathway). Our data also suggest that specific DNA structures may act act as important intermediates for [*MPH1*^+^] phenotypes. The linking of diverse physiological outcomes to a single epigenetic state suggests that [*MPH1*^+^] is a coordinated program that fuels specific and heritable changes in DNA repair networks. During periods of nucleotide starvation, when cells are ill-suited to their environments, they can acquire [*MPH1*^+^] at higher rates. This improves their chances of survival and enhances phenotypic diversification of the next generation. Mph1 is not alone in its capacity to assemble in response to replication fork stress (Xue et al., 2014). Many DNA repair factors localize to large assemblies to exert their functions. Our data provide an example in which such an assembly can encode a new set of activities that are heritable over long biological timescales, with the capacity to permanently hardwire adaptive phenotypic diversity into the genome.

## Acknowledgements

This work was supported by a National Institutes of Health New Innovator Award (NIH-DP2-GM119140), an NSF career award (NSFMCB1453762), a Searle Scholar Award (14-SSP-210), a Kimmel Scholar Award (SKF-15-154), and by a Science and Engineering Fellowship from the David and Lucile Packard Foundation to DFJ. JSB was supported by National Institutes of Health Training Grant (5T32GM007790-36). DMG was supported by a postdoctoral fellowship from the NIH (F32-GM109680). We thank members of the Jarosz laboratory as well as K. Cimprich (Stanford) and X. Zhao (MSKCC) for helpful comments, and S. Larios (Stanford) for reagent support.

## FIGURE LEGENDS

**Supplemental Figure 1.**
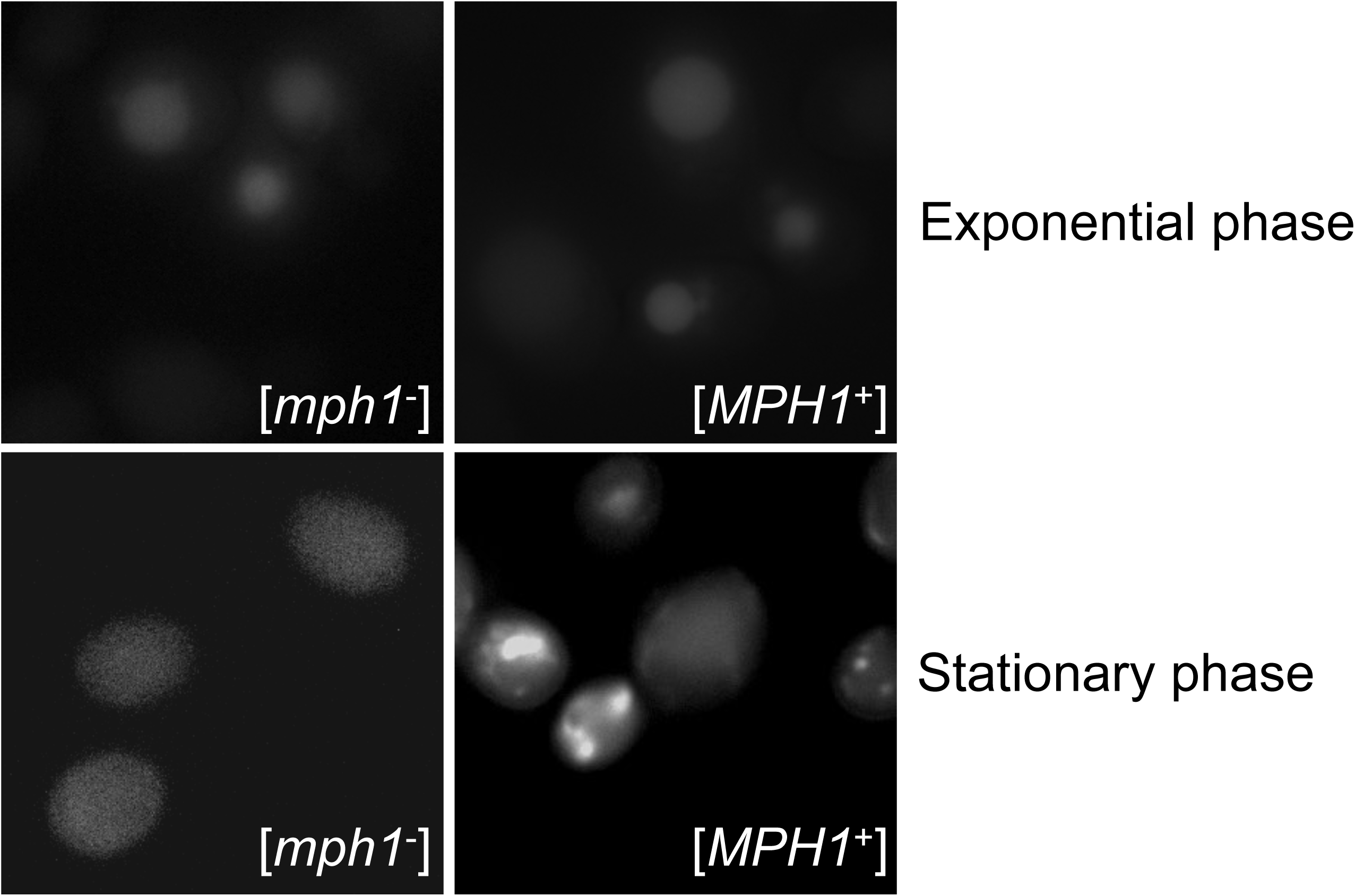
Localization of Mph1 protein in [*MPH1*^+^] cells. Fluorescence micrographs from diploid cells harboring [*MPH1*^+^], or [*mph1*^-^] control, each expressing integrated *MPH1-YFP* from its endogenous promoter. Exponential phased cells were imaged at OD_600_ ~0.5-0.8, stationary phase at OD_600_ ~1.5-2.

**Supplemental Figure 2.**
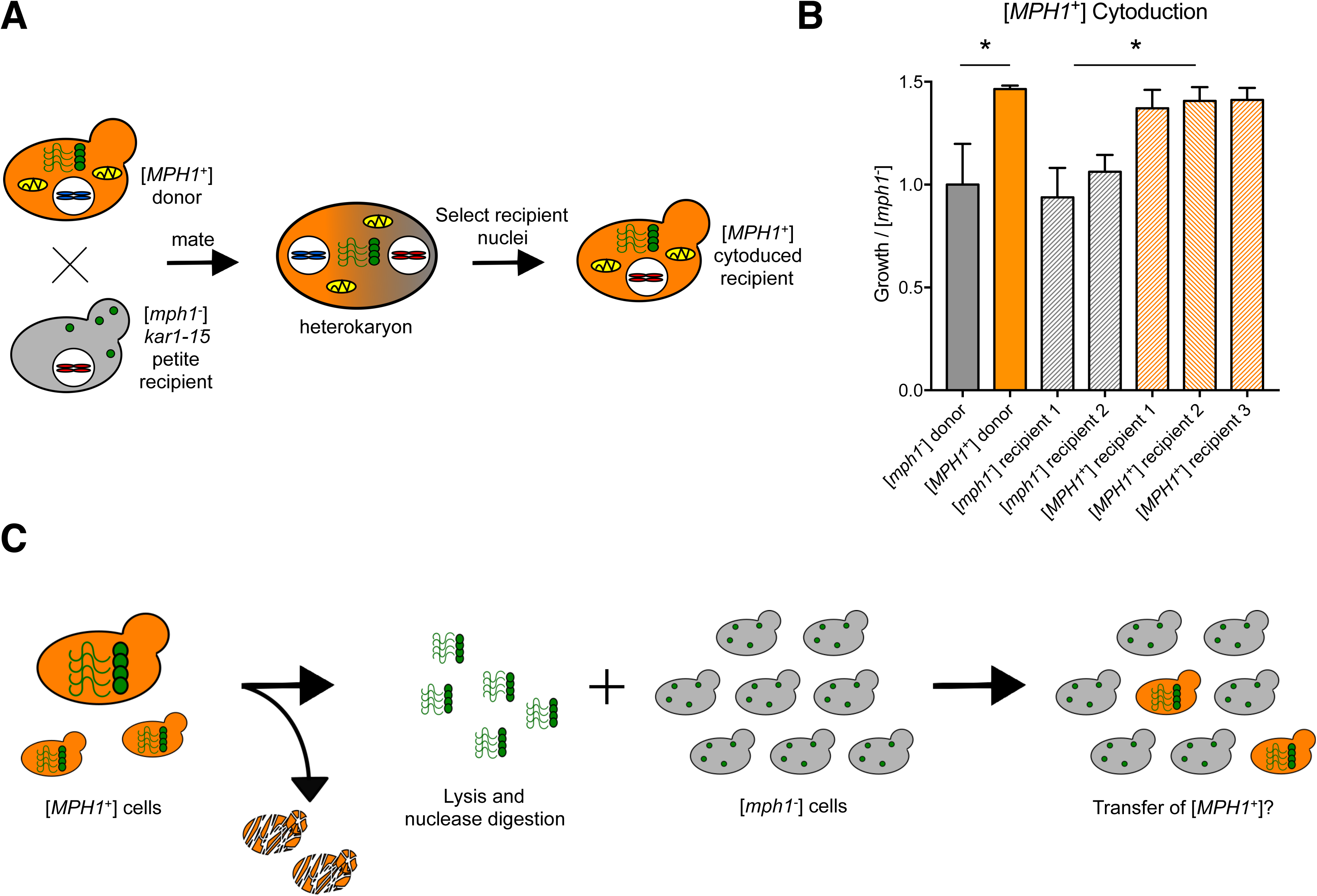
[*MPH1*^+^] is a cytoplasmic element. (A). Experimental schema of cytoduction experiments. (B) Growth of [*mph1*^-^] and [*MPH1*^+^] cytoduced strains in a DNA stressor to which [*MPH1*^+^] promotes resistance (10 μM mycophenolic acid). Error bars represent SEM from 3 biological replicates. (C) Experimental schema for protein transformation experiments.

**Supplemental Figure 3.**
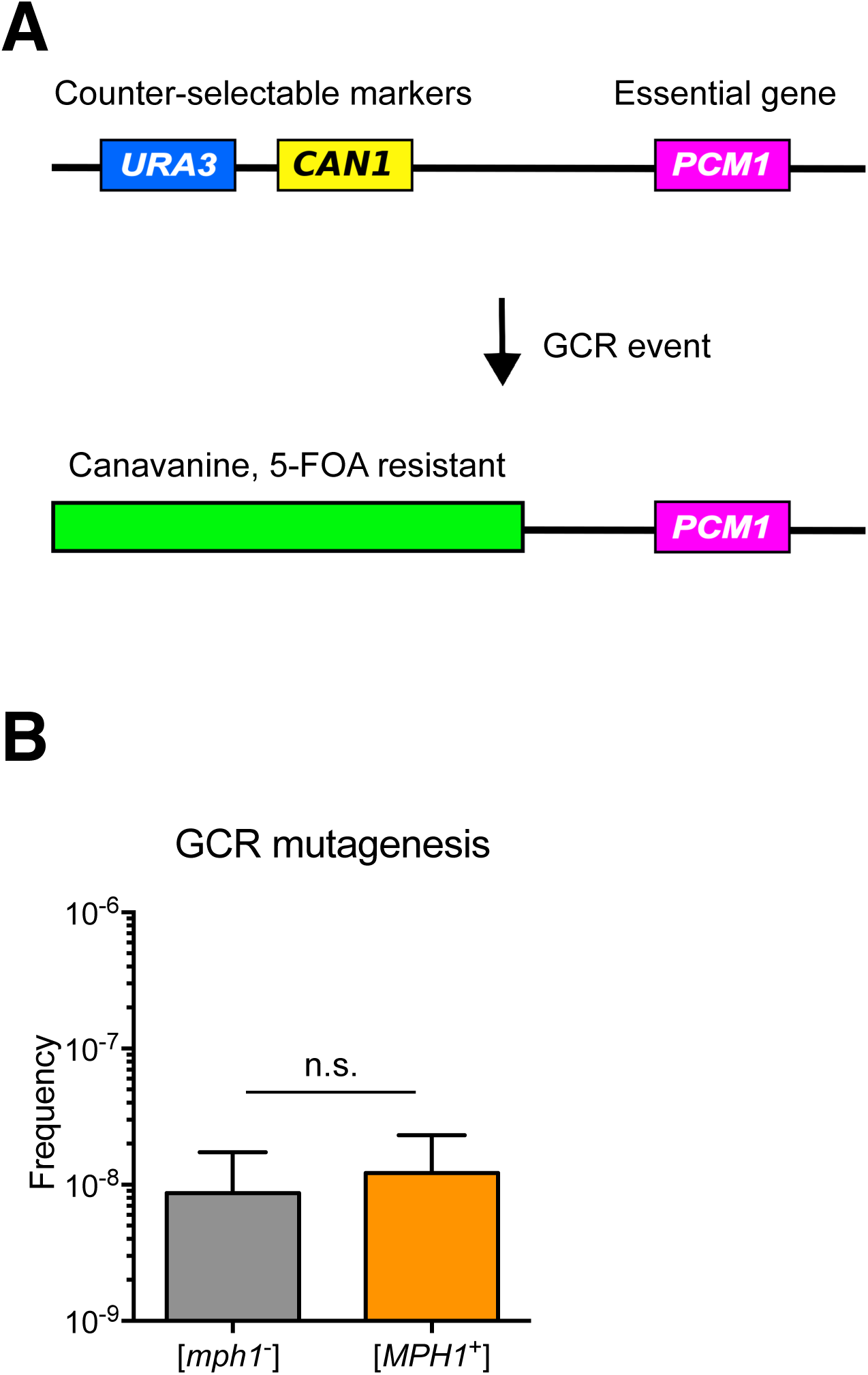
[*MPH1*^+^] does not influence GCR mutagenesis. (A) Diagram of GCR reporter used. Two counter-selectable markers (*URA3* and *CAN1*) are located upstream of an essential gene (*PCM1*). When cells are presented with a double counter-selection, the probability of mutating both genes in a single generation is vanishingly small. Therefore, resistant colonies will undergo a GCR event at the precise location, preserving the essential gene. (B) GCR frequencies in [*mph1*^-^] and [*MPH1*^+^] strains. Error bars represent SEM from 5 biological replicates.

**Supplemental Figure 4.**
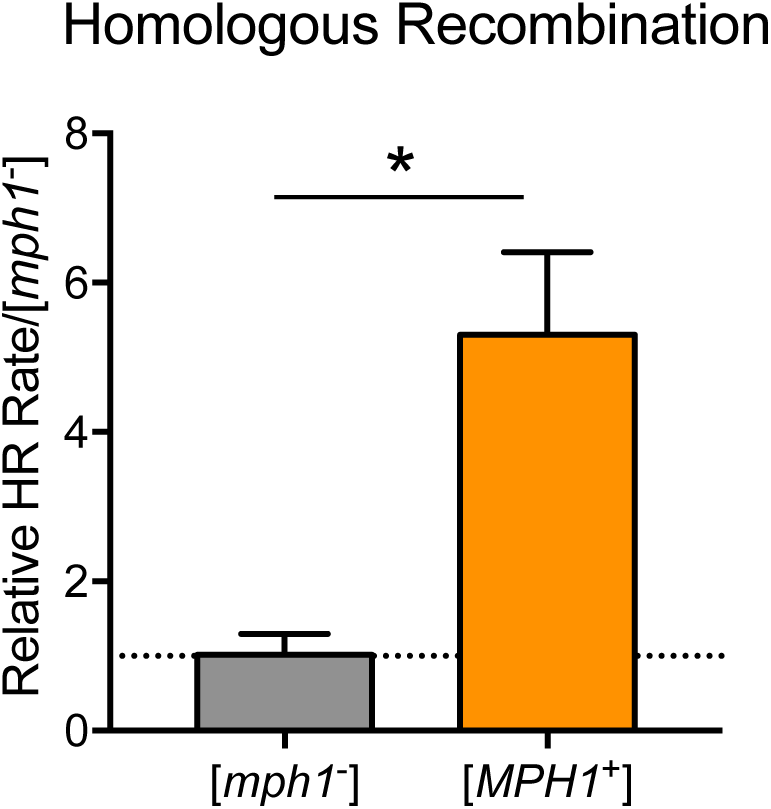
[*MPH1*^+^] increases homologous recombination. Normalized genomic integration frequencies of a linear DNA cassette in [*mph1*^-^] and [*MPH1*^+^] strains. Error bars represent SEM for 6 biological replicates.

**Supplemental Figure 5.**
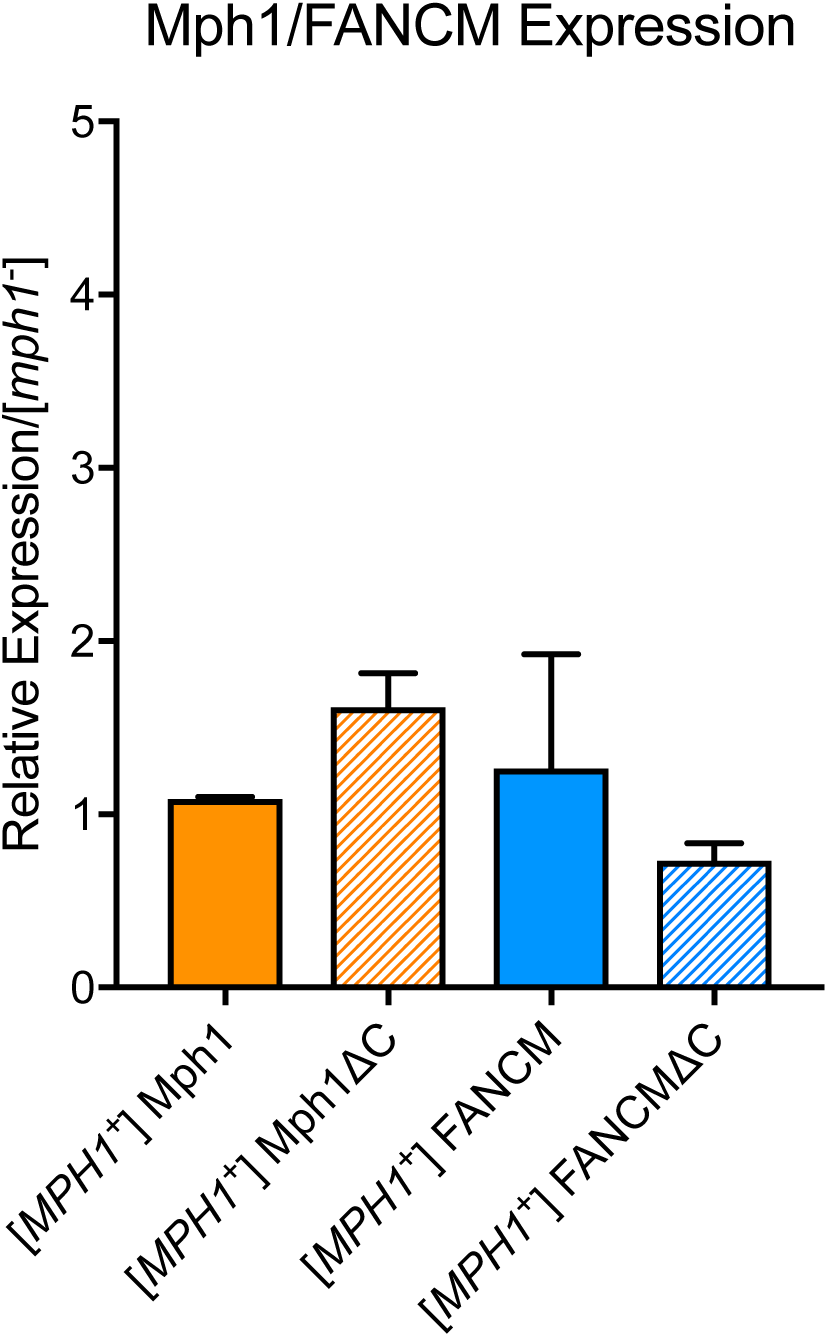
Expression levels of FANCM and Mph1 variants. Bar graph showing the relative expression levels of FANCM and Mph1 plasmids in [*MPH1*^+^] vs. [*mph1*] strains measured using RT-PCR. Error bars represent SEM from 3 biological replicates.

**Supplemental Figure 6.**
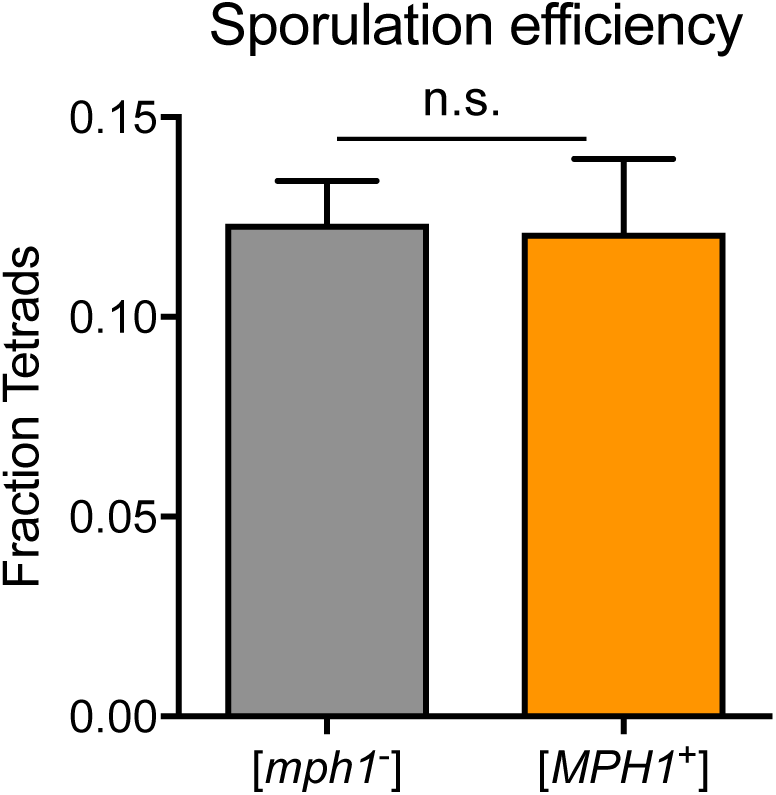
[*MPH1*^+^] does not affect sporulation. Fraction of tetrads in [*mph1*^-^] and [*MPH1*^+^] strains after 5 days. Error bars represent SEM from 3 biological replicates.

**Supplemental Figure 7.**
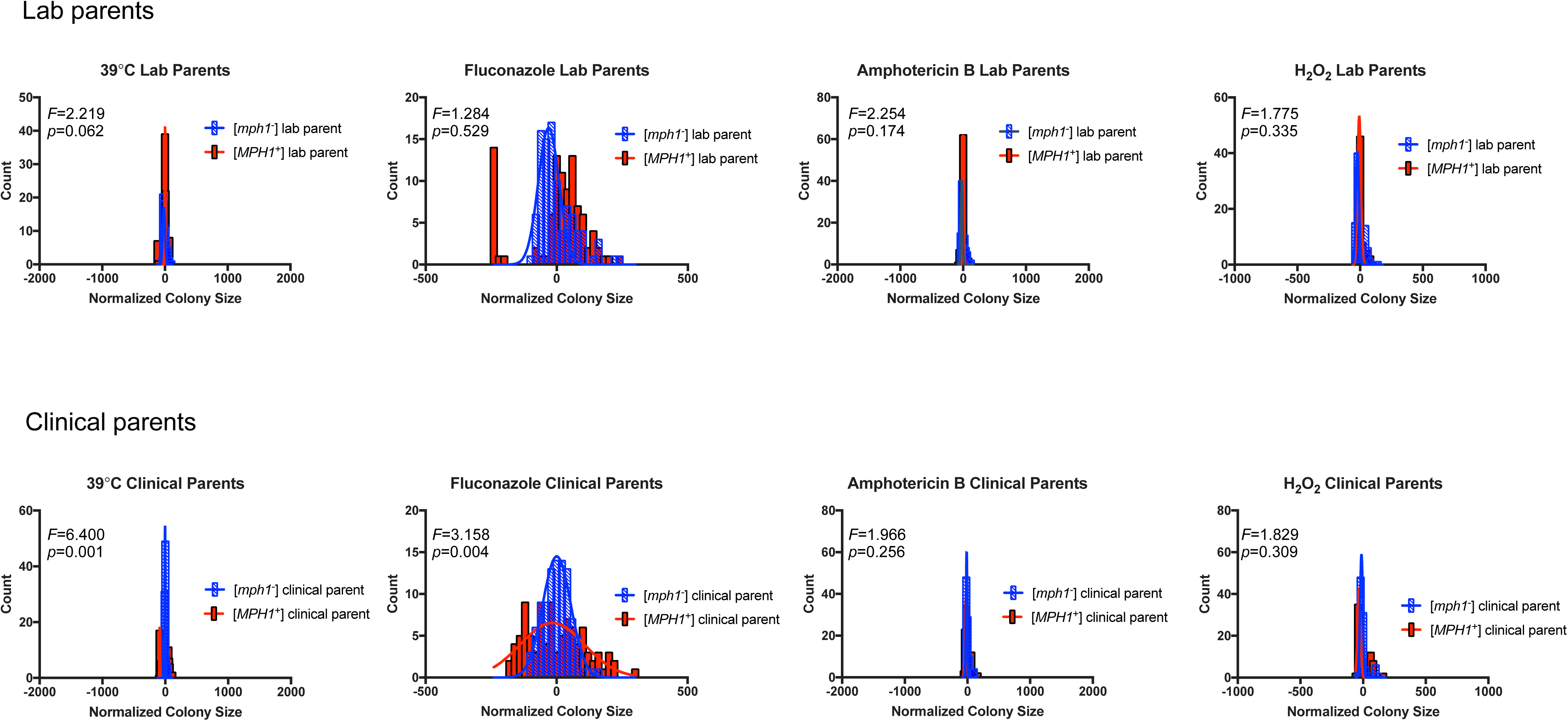
Phenotypic variation in [*mph1*^-^] and [*MPH1*^+^] derivatives of laboratory and clinical parent strains. Phenotypic distributions of [*mph1*^-^] or [*MPH1*^+^] parental strains for wild cross in the 4 different stressors. Range of x-axis used is identical to each corresponding progeny histogram in Figure 5.

**Supplemental Figure 8.**
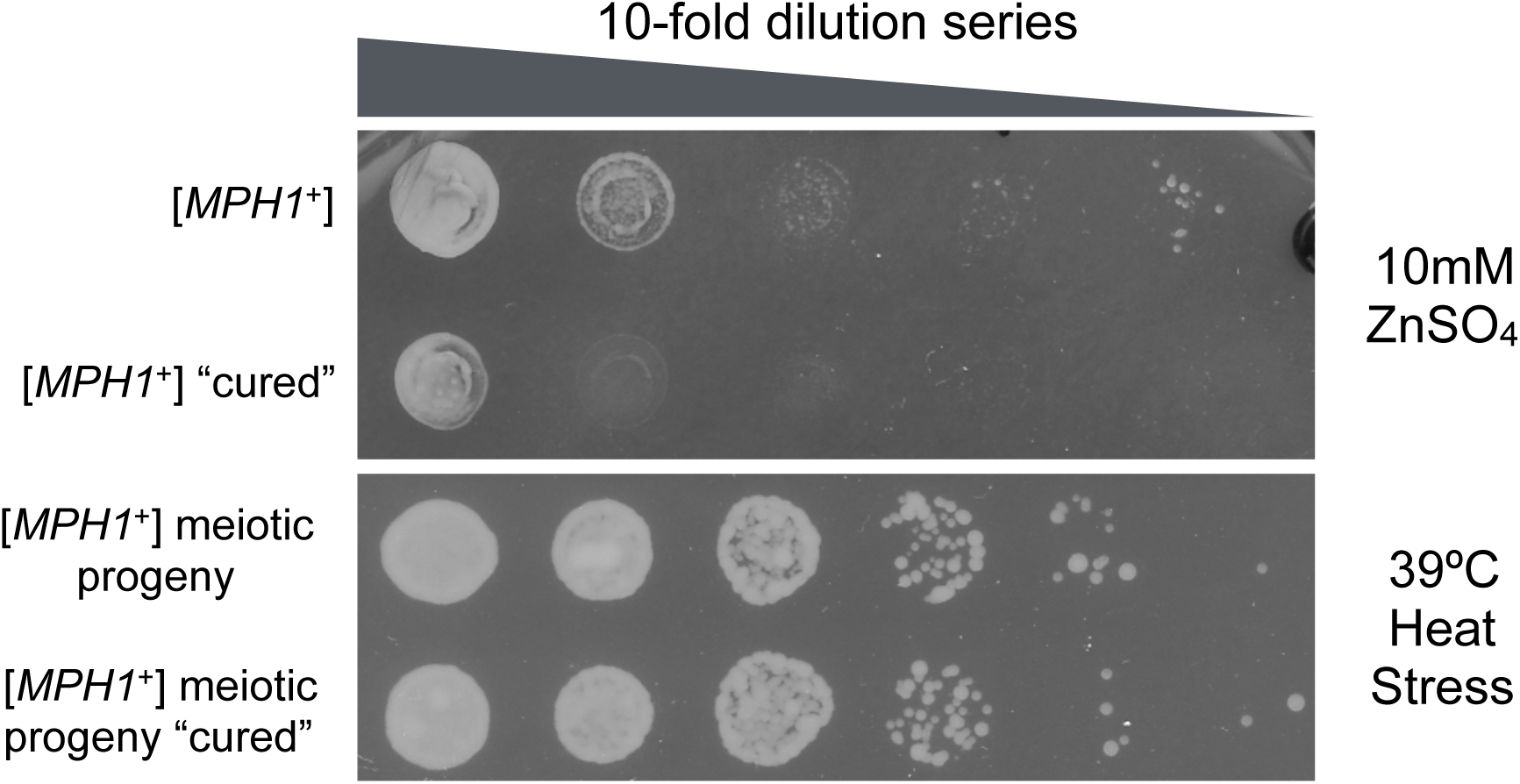
Mph1 does not act as a phenotypic capacitor. Top panel: [*MPH1*^+^] parent strain before and after prion ‘curing’ spotted in a 10-fold dilution series on a plate containing 10 mM ZnSO_4_. Bottom panel: Heat-resistant progeny from wild outcross with an [*MPH1*^+^] parent before and after prion ‘curing’ spotted in a 10-fold dilution series on a YPD plate grown at 39 °C.

**Supplemental Table 1.** Yeast strains used in this study.

| Strain | Source |
| --- | --- |
| *mph1-Q603D-3Flag::HIS3* | Xiaolan Zhao lab (MSKCC) |
| *BY4742 rad5Δ* | Yeast MATα deletion library |
| *BY4742 rad51Δ* | Yeast MATα deletion library |
| *BY4742 sgs1Δ* | Yeast MATα deletion library |
| *BY4742 srs2Δ* | Yeast MATα deletion library |
| *BY4742 mhf1Δ* | Yeast MATα deletion library |
| *BY4742 mhf2Δ* | Yeast MATα deletion library |
| *BY4742 chl1Δ* | Yeast MATα deletion library |
| *BY4742 exo1Δ* | Yeast MATα deletion library |
| *BY4742 mgm101Δ* | Yeast MATα deletion library |
| *BY4742 msh2Δ* | Yeast MATα deletion library |
| *BY4742 msh6Δ* | Yeast MATα deletion library |
| *BY4742 pso2Δ* | Yeast MATα deletion library |
| *BY4742 slx4Δ* | Yeast MATα deletion library |
| *BY4741 mph1Δ* | Yeast MATα deletion library |
| YJM975 | SGRP collection |
| Mph1-YFP | Xiaolan Zhao lab (MSKCC) |
| *MDG1::K.lactisURA3* prion reporter | Chakrabortee et al., 2016 |

**Supplemental Table 2.** Plasmids used in this study.

| Plasmid | Source |
| --- | --- |
| pUG72 | Euroscarf |
| 416-GPD-ccdB-eGFP | Addgene Yeast Gateway kit |
| 416-GPD-Mph1-eGFP | Addgene Yeast Gateway kit; Yeast ORFeome collection E. coli (YSC3868) |
| 416-GPD-Mph1ΔC-eGFP | Addgene Yeast Gateway kit; Yeast ORFeome collection E. coli (YSC3868) |
| 416-GPD-FANCM-eGFP | Addgene Yeast Gateway kit; FANCM from Stephen West lab |
| 416-GPD-FANCMΔC-eGFP | Addgene Yeast Gateway kit; GE Life Sciences Human ORFeome library |
| Hsp70(K69M) | Jarosz et al., 2014b |

**Supplemental Table 3.**
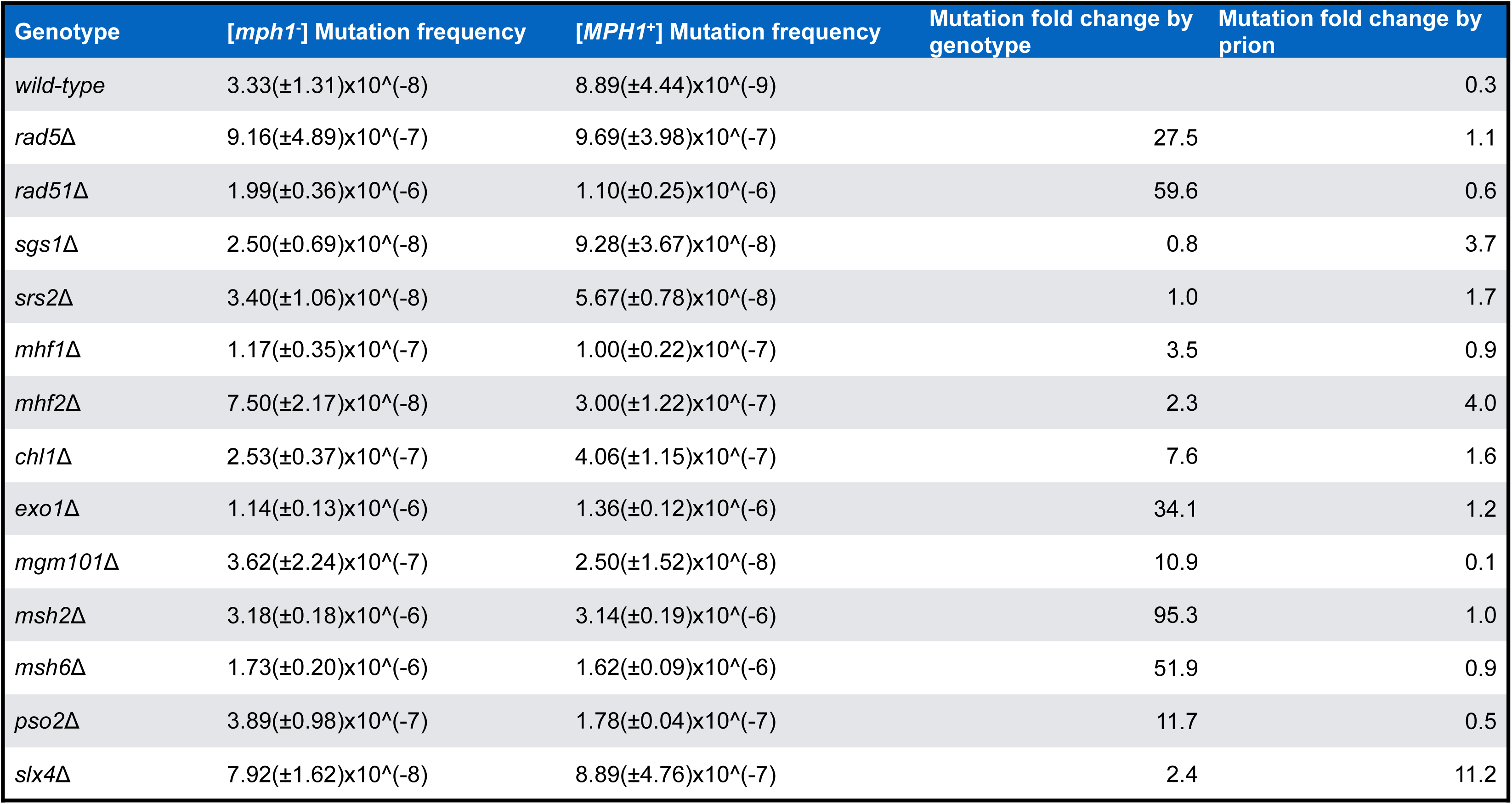
Interacting proteins required for [*MPH1*^+^]-dependent anti-mutator phenotypes. Table showing [*mph1*^-^] and [*MPH1*^+^] mutation frequencies for haploid progeny harboring genetic knockouts of DNA repair factors. 1^st^ column – genotypes of each strain assayed; 2^nd^ column – mutation frequency with SEM for each knockout in a naïve [*mph1*^-^] strain; 3^rd^ column – mutation frequency with SEM for corresponding [*MPH1*^+^] strain; 4^th^ column – Fold-change in mutation frequency by each genetic knockout compared to a wild-type naïve [*mph1*^-^] strain; 5^th^ column – Fold-change in mutation frequency imparted by [*MPH1*^+^] for the same genotype.

| Genotype | [*mph1*<sup>-</sup>] Mutation frequency | [*MPH1*<sup>+</sup>] Mutation frequency | Mutation fold change by genotype | Mutation fold change by prion |
| --- | --- | --- | --- | --- |
| *wild-type* | 3.33(±1.31)x10<sup>^</sup>(-8) | 8.89(±4.44)x10<sup>^</sup>(-9) |  | 0.3 |
| *rad5Δ* | 9.16(±4.89)x10<sup>^</sup>(-7) | 9.69(±3.98)x10<sup>^</sup>(-7) | 27.5 | 1.1 |
| *rad51Δ* | 1.99(±0.36)x10<sup>^</sup>(-6) | 1.10(±0.25)x10<sup>^</sup>(-6) | 59.6 | 0.6 |
| *sgs1Δ* | 2.50(±0.69)x10<sup>^</sup>(-8) | 9.28(±3.67)x10<sup>^</sup>(-8) | 0.8 | 3.7 |
| *srs2Δ* | 3.40(±1.06)x10<sup>^</sup>(-8) | 5.67(±0.78)x10<sup>^</sup>(-8) | 1.0 | 1.7 |
| *mhf1Δ* | 1.17(±0.35)x10<sup>^</sup>(-7) | 1.00(±0.22)x10<sup>^</sup>(-7) | 3.5 | 0.9 |
| *mhf2Δ* | 7.50(±2.17)x10<sup>^</sup>(-8) | 3.00(±1.22)x10<sup>^</sup>(-7) | 2.3 | 4.0 |
| *chl1Δ* | 2.53(±0.37)x10<sup>^</sup>(-7) | 4.06(±1.15)x10<sup>^</sup>(-7) | 7.6 | 1.6 |
| *exo1Δ* | 1.14(±0.13)x10<sup>^</sup>(-6) | 1.36(±0.12)x10<sup>^</sup>(-6) | 34.1 | 1.2 |
| *mgm101Δ* | 3.62(±2.24)x10<sup>^</sup>(-7) | 2.50(±1.52)x10<sup>^</sup>(-8) | 10.9 | 0.1 |
| *msh2Δ* | 3.18(±0.18)x10<sup>^</sup>(-6) | 3.14(±0.19)x10<sup>^</sup>(-6) | 95.3 | 1.0 |
| *msh6Δ* | 1.73(±0.20)x10<sup>^</sup>(-6) | 1.62(±0.09)x10<sup>^</sup>(-6) | 51.9 | 0.9 |
| *pso2Δ* | 3.89(±0.98)x10<sup>^</sup>(-7) | 1.78(±0.04)x10<sup>^</sup>(-7) | 11.7 | 0.5 |
| *slx4Δ* | 7.92(±1.62)x10<sup>^</sup>(-8) | 8.89(±4.76)x10<sup>^</sup>(-7) | 2.4 | 11.2 |

